# In vivo efficacy of L-ascorbic acid in restricting cholera pathogenesis

**DOI:** 10.1101/2025.04.01.646656

**Authors:** Prolay Halder, Himanshu Sen, Sanjib Das, Pritam Nandy, Asish Kumar Mukhopadhyay, Jeffrey H. Withey, Saumya Raychaudhuri, Hemanta Koley

**Author notes:** Corresponding authors: Jeffrey H. Withey, Professor, Department of Biochemistry, Microbiology, and Immunology, Wayne State University School of Medicine, Detroit, MI, United States, Hemanta Koley, Scientist F, Division of Bacteriology, ICMR-National Institute for Research in Bacterial Infections, P-33, CIT Road, Scheme-XM, Beliaghata, Kolkata-700010, India, /.

## Abstract

In the last three centuries, our world has experienced seven cholera pandemics. Cholera is a deadly diarrheal disease caused by *Vibrio cholerae*, an important member of the gamma-proteobacteria. Despite conventional treatments like antibiotics, vaccines, and ORS, the 7th cholera pandemic is still a major threat to developing nations. L-ascorbic acid was recently shown to effectively kill the *Vibrio cholerae* cells *in vitro* under various growth conditions mimicking the *in vivo* host conditions, including growth in the presence of bile salts, growth of acid-adapted V. cholerae, and growth in the presence of various ORS components. In the current study, we extend this work and test the efficacy of L-ascorbic acid in animal models (rabbit ileal loop and the removable intestinal tie adult rabbit diarrhea). We show that L-ascorbic acid can effectively reduce the bacterial load in both the rabbit models as well as fast-track the recovery from the diarrheal symptoms.

**Importance:** Cholera is a severe human diarrheal disease that affects millions each year. Cholera treatment is primarily oral rehydration to balance the voluminous fluid loss. Previous studies have found that L-ascorbic acid (L-AA) is effective at inhibiting growth of Vibrio cholerae, the causative agent of cholera. Thus L-AA has potential as am inexpensive cholera therapeutic. Here, L-AA is tested in rabbit models to determine whether it can inhibit the effects of V. cholerae infection. Results suggest that L-AA treatment inhibits bacterial proliferation and leads to shortened recovery from disease.

## Introduction

Cholera continues to ravage the world. A recent global analysis of cholera cases strongly advocates the upsurge of cholera. Since the 19^th^ century, cholera has been studied intensively in various fields, from basic science to therapeutics^1^. Several prevention and intervention strategies are now practiced worldwide to combat and control cholera^2^. Other than mainstays of conventional treatments (e.g., oral rehydration therapy, antibiotics, and vaccines), new interventions such as probiotics and phage therapy have emerged as promising approaches to tackle cholera infections^3-5^. Recently, L-ascorbic acid (L-AA) has been demonstrated to mitigate effectively the viability of various clinical *Vibrio cholerae* strains under various growth conditions. Further analysis also revealed its efficacy in reducing bile-induced biofilm development of such strains^6^. In the present work, we wanted to examine the anti-*Vibrio cholerae* activity of L-AA in a mammalian infection model of cholera. Towards this end, rabbit ileal loop (RIL) and the removable intestinal tie adult rabbit diarrhea (RITARD) models were selected in the present work. Our data demonstrated that L-ascorbic acid (L-AA) could effectively reduce fluid volume and bacterial burden in stools in both RIL and RITARD animal models, respectively.

## Materials and methods

### Bacterial culture

Overnight grown culture of *Vibrio cholerae* strain N16961 (O1, El Tor, Ogawa, streptomycin resistant) was diluted in Luria Bertani broth and grown to mid log phase at 37° C with shaking. This culture was then used for the rabbit infections.

### Animal experiments

A rabbit ileal loop (RIL) experiment was used to assess the total pathogenicity potential of *V. cholerae* strain N16961^7^. The rabbits were put under anesthesia, followed by a short surgical incision to expose the intestine. A maximum of six ligated ileal loops (each measuring about 10 cm in length) were produced in each rabbit. The culture suspension containing approximately 10^9^ colony forming units (CFUs) or normal saline was administered to each ligated intestinal loop in a volume of 1 mL. In order to ensure the rabbits’ survival, the ligated intestine was then replaced and sutured. Following an 18-hour challenge, the rabbits were euthanized, and the fluid accumulation (FA) ratio was recovered from the loops and quantified. The mean volume (mL) of fluid accumulated per unit length (cm) of each loop was used to express the results obtained from triplicate experiments. An increase in the FA ratio was correlated with an increase in virulence, and vice versa. Based on FA ratios of ≤ 0.2 and ≥ 1.0, respectively, negative and strong positive reactions were evaluated.

The RITARD model was run using Roy et al.’s methodology^8^. Following a 24 h withholding of food, the animals were sedated using anesthesia, and the cecum was exposed surgically. A cotton thread was used to permanently knot the caecum 1.2 cm away from the ileocecal junction without rupturing any nearby blood supply. *Vibrio cholerae* N16961 alone or with L-AA (600 mg) was injected into the lumen and a detachable slip-knot tie was placed on the ileum. The incision was closed after placing the colon and cecum back into the peritoneal cavity. One end of the tie was kept free outside in a neatly tucked condition. After the rabbits regained consciousness, the reversible tie was loosened two hours post infection, and rabbits were maintained on sterilized water and chow. The clinical indicators of diarrhea, the stool shedding, and other physiological symptoms were monitored in rabbits for next two days, at 24 h intervals. Every day, fresh feces and rectal swabs were collected to track shedding. Stool samples were serially diluted and plated onto Luria Bertani agar plates containing streptomycin (100 µg/mL) to quantify the shedding of the challenge bacterium. The colonies were further verified using specified primers in PCR based method and a suitable antiserum for slide agglutination.

## Results

To determine the effect of L-ascorbic acid (L-AA) on cholera toxin activity, the rabbit ileal loop RIL model was used. This model enables quantification of fluid excreted due to *V. cholerae* infection. L-AA had a considerable effect on *Vibrio cholerae* N16961, as demonstrated using the RIL (**Figure 1A**). After 18 h of infection, a notable reduction in fluid volume was observed in the loop containing L-AA (600 mg) with *Vibrio cholerae* N16961 (10^9^ cells) as compared to the positive control without L-AA (**Table 1**). This observation remained consistent when the L-AA concentration was reduced to 150 mg. CFUs recovered from the loops further supported the fluid accumulation results, indicating that L-AA inhibited *V. cholerae* growth and/or colonization (**Table 1**). One explanation for this is that L-AA’s high concentration and extremely acidic pH might have a negative impact on bacterial viability^6^. Although the fluid accumulation was reduced, bacterial CFUs remained unchanged at the lower concentration of L-AA (150 mg) indicating a bacteriostatic effect on bacterial viability (**Figure 1B**).

**Table 1.** Description of the loops made for RILA model.

| Loop no. | Loop description | Mean loop length (cm) | Standard deviation | Mean accumulated fluid volume (ml) | Standard deviation |
| --- | --- | --- | --- | --- | --- |
| 1 | PBS only (2ml) | 11.16666 | ±0.6236 | 0 | ±0 |
| 2 | L-AA 150 mg | 10.66666 | ±0.9428 | 0 | ±0 |
| 3 | L-AA 600 mg | 10.66666 | ±0.2357 | 1.66666 | ±1.24721 |
| 4 | L-AA 150 mg + N16961 (10<sup>9</sup> cells) | 10.83333 | ±0.4714 | 13 | ±1.63299 |
| 5 | L-AA 600 mg + N16961 (10<sup>9</sup> cells) | 10.33333 | ±0.4714 | 9.16666 | ±0.62361 |
| 6 | N16961 (10<sup>9</sup> cells) | 10.16666 | ±0.2357 | 23.5 | ±1.22474 |

**Figure. 1:**
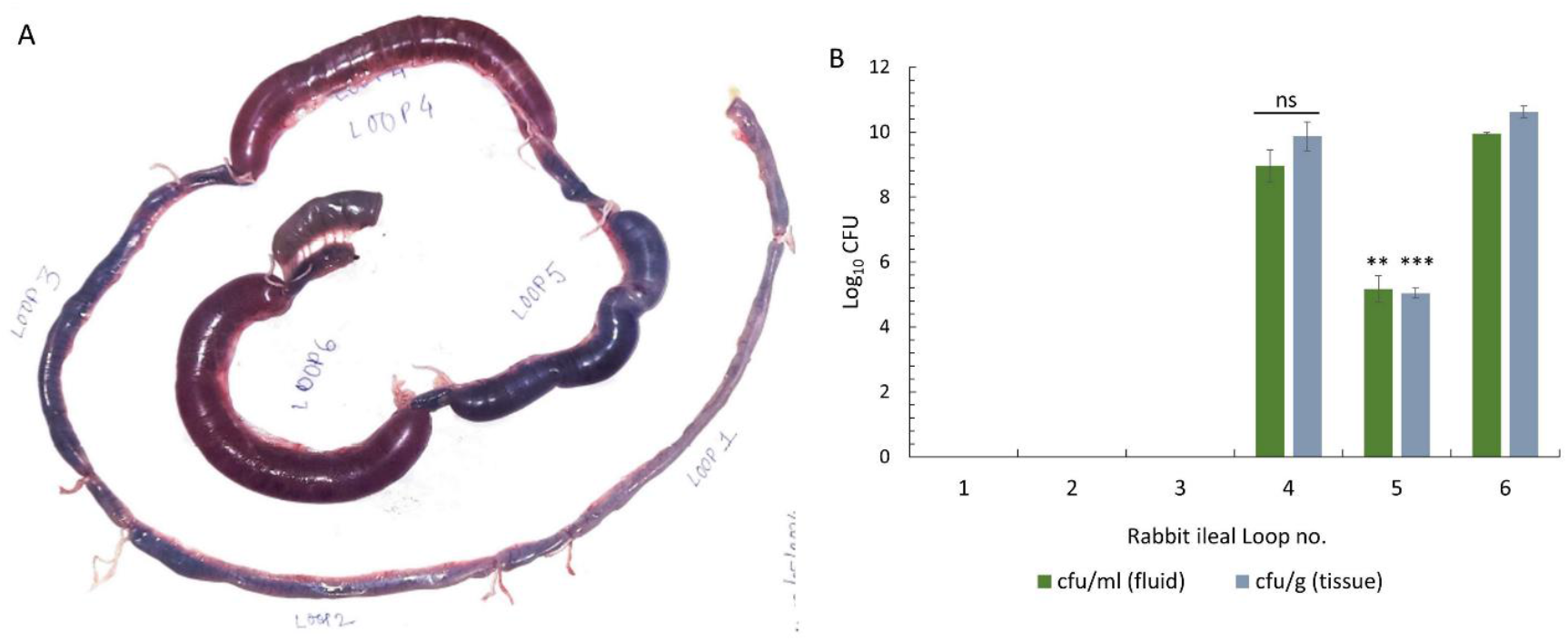
Rabbit ileal Loop (RIL) assay. **(A)** RIL response with culture of N16961, N16961 + L-ascorbic acid (600 mg/2ml), N16961 + L-ascorbic acid (150 mg/2ml), L-ascorbic acid (600 mg/2ml), L-ascorbic acid (150 mg/2ml) and PBS as negative control. Image shown is a representative of the experiment done in triplicate. **(B)** Bacterial count in fluid (CFU/ml) and tissue section (CFU/gm) of different loops of RIL assay. Bar graph was plotted as the mean of the CFU/ml and CFU/gm of tissue from the values of the biological triplicate data. Error bars represent the ±standard deviation. (ns = data not significant; ** = *p value* 0.01 < > 0.001; *** = *p value* < 0.001).

To further assess the possible impact of L-AA on *Vibrio cholerae* infection, the removable intestinal tie-adult rabbit diarrhea (RITARD) model was used. In the RITARD experiments, all the rabbits received *V. cholerae* either with or without any L-AA water diet. Rabbits that received 600 mg of L-ascorbic acid along with the *V. cholerae* infection did not exhibit any symptoms of infection. Following surgery, L-AA was given as a supplement to all rabbits except the positive control group. The rabbits with a regular water diet the day before the surgery had clear signs of diarrhea, including soft, loose blobs of stool and severe physical weakness. In addition to the positive control (N16961-infected, no L-AA) rabbit, loose stool (soft, loose blobs) and diarrhea symptoms were also observed in the N16961-infected rabbit given an L-AA (300 mg/100 ml) supplemented water diet (**Figure 2A**). In contrast, the N16961-infected rabbit that received a 600 mg dose of L-AA had no symptoms of diarrhea and a normal stool composition. Stool pellets became regular, tiny, spherical, and black in color. On the second day of infection, rabbit + N16961, which was fed a regular water diet without L-AA, had acute physical debility and diarrhea (**Figure 2A**). Unfortunately, the rabbit was found dead later that day. Remarkably, the N16961-infected rabbit treated with L-AA (300 mg/100 ml) supplemented water diet also returned to normal stool consistency, whereas the L-AA (600 mg) inoculated N16961-infected rabbit treated with L-AA (300 mg/100 ml) water diet had normal stool consistency. All the animals receiving L-AA treatment had regular, small, spherical, and normal-shaped feces pellets on the second day of infection; they were also entirely black in color, much like an uninfected rabbit (**Figure 2A**).

**Figure. 2:**
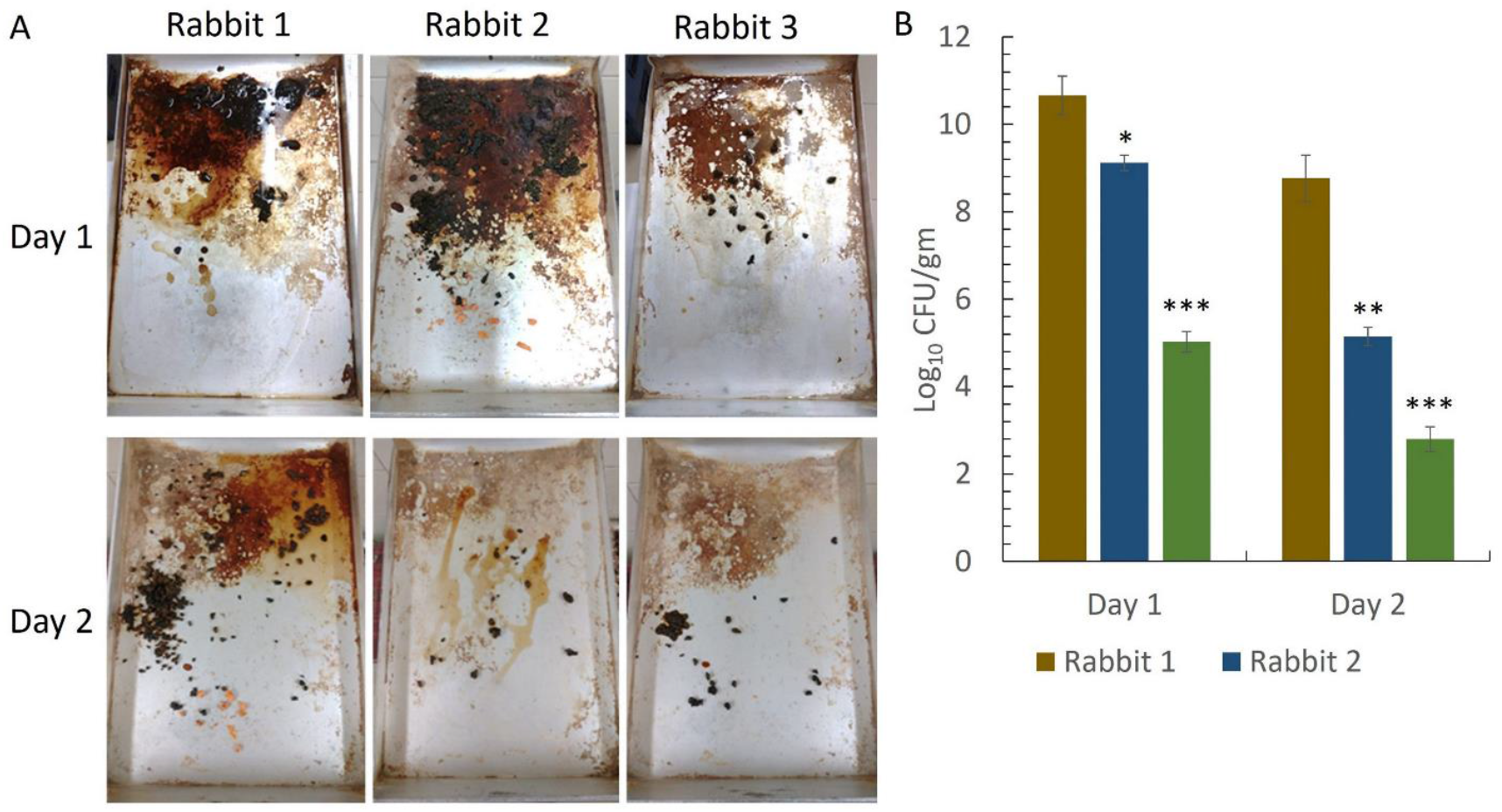
Protection study by RITARD model. **(A)** Stool pathology after *Vibrio cholerae* (N16961) infection in RITARD model in rabbits. Rabbit 1: showed a distinct diarrheal symptoms on day 1 and day 2 after surgery. Rabbit 2: showed diarrheal symptoms on day 1 and recovered in normal stool consistence on day 2 after surgery. Rabbit 3: there were no diarrheal symptoms on days 1 and 2 after surgery, **(B)** Bacterial shedding in stool (CFU/gm) of different groups of rabbits after RITARD assay. Bacterial count was very low in rabbit 3 and rabbit 2 in comparison to rabbit 1 for both day 1 and day 2. [Rabbit 1: N16961 infected (regular water diet), rabbit 2: N16961-infected (L-ascorbic acid water diet), rabbit 3: N16961 (inoculated in L-ascorbic acid dissolved water) infected and L-ascorbic acid water diet.]. Bar graph was plotted as the mean of the CFU/gm of tissue from the values of the biological triplicate data. Error bars represent the ±standard deviation. (* = *p value* 0.05 < > 0.01; ** = *p value* 0.01 < > 0.001; *** = *p value* < 0.001).

In the RITARD assay, we also saw a clear distinction between the bacterial secretion in the stool of rabbits treated with L-AA or untreated with L-AA (**Figure 2B**). We further conclude from this RITARD model that L-AA plays a significant role in the reduction, elimination, and recovery from *in vivo V. cholerae* infections.

## Discussion

It is important to recognize that cholera infection causes tremendous shifts in the gut microbiota. Therefore, it is a challenge to reestablish the gut community structure while recovering from cholera. Vitamin C has been recognized to exert a positive influence on gut microbiota^9^. Recently, *E. coli* Nissle 1917 (EcN 1917), in combination with prebiotic sugars, has been evidenced to modulate the community structure of colonic microbiota in a small-scale host-free microbiota system^10^. The present work using in vivo animal models reconfirms a previous study on the effectiveness of L-AA *in vitro* in a dose-dependent manner^6^. Thus, LAA holds excellent potential in reducing the number of diarrhea cases due to *V. cholerae* in endemic areas. Incorporation of L-AA to oral rehydration solution (ORS) might revolutionize existing methods of rehydration treatment.

It should be noted that *EcN* 1917, along with glucose, is efficacious in restricting the growth of *V. cholerae* both *in vitro* and in zebrafish infection models^3,4^. It would be interesting to examine the effectiveness of a consortium of L-AA, probiotic, and prebiotic sugars to not only mitigate the pathogenesis of *V. cholerae* but also aid in shaping the gut community structure. Additional studies are necessary to address this issue.

